# Partial Unwinding of Transmembrane Helices Facilitates Alternating Access in the Neurotransmitter:Sodium Symporter LeuT

**DOI:** 10.1101/156968

**Authors:** Patrick S. Merkle, Kamil Gotfryd, Michel A. Cuendet, Katrine Z. Leth-Espensen, Ulrik Gether, Claus J. Loland, Kasper D. Rand

## Abstract

The prokaryotic neurotransmitter:sodium symporter (NSS) LeuT from *Aquifex aeolicus* is an established structural model for mammalian NSS counterparts. Here, we investigate the substrate translocation mechanism of LeuT by measuring the solution-phase structural dynamics of the transporter in distinct functional states by hydrogen/deuterium exchange mass spectrometry (HDX-MS). Our HDX-MS data pinpoint LeuT segments involved in substrate transport and reveal for the first time a comprehensive and detailed view of the solution-phase dynamics associated with transition of the transporter between outward- and inward-facing configurations in a Na^+^- as well as K^+^-dependent manner. The results suggest that partial unwinding of transmembrane helices 1/5/6/7 drive LeuT from a substrate-bound, outward-facing occluded conformation towards an inward-facing open state. We thereby envisage that substrate release is facilitated by formation of two distinct solvent pathways, which mediate access to the Na^+^ and substrate binding sites.

The neurotransmitter:sodium symporter (NSS) family includes prokaryotic and eukaryotic integral membrane proteins that harness the energy stored in the Na^+^ concentration gradient to transport solutes across the cell membrane (1, 2). Mammalian NSS proteins play an important role in terminating the neurotransmission in the central nervous system by rapid uptake of neurotransmitters against their concentration gradient into the presynaptic neuron or neighboring glial cells (3-5). Numerous neuropsychiatric conditions are associated with altered function of NSS transporters or low neurotransmitter concentrations in the synaptic cleft (2). Many NSS proteins therefore represent favorable targets for therapeutic drugs that act as potent transport inhibitors to increase neurotransmitter levels at the synaptic junction (2, 6). Despite the importance of mammalian NSS proteins in neurobiology and pharmacology, the molecular mechanisms underlying the transport function of mammalian NSS family members have proven difficult to elucidate by traditional methods as the target proteins are unstable in detergent-solubilized form (7) and difficult to purify in required quantity and purity.

The prokaryotic NSS protein LeuT from *Aquifex aeolicus* has emerged as an important structural model to investigate the structure-function relationship of mammalian NSS counterparts (7-9). High-resolution crystal structures of LeuT in distinct functional states (10-13) have established a structural framework and given rise to mechanistic models depicting the substrate transport mechanism (14, 15). LeuT comprises 12 transmembrane segments (TMs) that are interconnected by relatively short extracellular (EL) and intracellular (IL) loops. The first TMs 1-5 share a similar structural repeat with the following TMs 6-10 but are inverted in the membrane plane (10). The specific arrangement of TM domains, also referred to as the ‘5+5’ or LeuT fold (10), has been observed in other NSS proteins (16-18), but also in transporters without sequence homology to LeuT (19, 20), suggesting the possibility of a conserved structural scaffold for many secondary active transporters (8). The primary binding site for the substrate (S1) and residues involved in coordination of two sodium ions (Na1 and Na2) are located approximately halfway across the membrane bilayer in the core of the transporter (10). The characteristic unwound regions of TMs 1 and 6 (providing both TMs with an a- and a b-section) fulfill a dual role in coordinating the sodium ions and forming interactions with the substrate molecule. Individual amino acid residues of TMs 3 and 8 as well as the sodium occupancy in the Na1 site complete the S1 binding pocket.

According to the widely embraced ‘alternating access’ model (21, 22), secondary active transporters isomerize between distinct functional states in a substrate-dependent manner. That is, the substrate binding site is alternatively exposed to either the intracellular or extracellular aqueous environment. X-ray crystallography provided structures of LeuT in ‘outward-facing open’ (11, 12), ‘outward-facing occluded’ (10), and ‘inward-facing open’ (11) conformations and led to the identification of external and internal gating residues in LeuT and related transporters (23-26). Based on these structural snapshots, it has been hypothesized that local and large-scale structural rearrangements are required to regulate the molecular gates and the outward-to-inward transition of the transporter, respectively (11). Combined evidence from crystallographic, functional, and simulation studies suggest that the underlying allosteric couplings are essential for LeuT to function as a symporter (27). Several key aspects of the transport cycle, in particular the molecular mechanism related to the transition of LeuT to the inward-facing open state, have remained controversial and are the subject of extensive debate (8, 28-30).

Here, we have studied the substrate translocation mechanism of LeuT by measuring the structural dynamics of the protein in solution as a function of time and substrate/ion composition (*e.g.* leucine, Na^+^, K^+^, and Cs^+^) by local hydrogen/deuterium exchange mass spectrometry (HDX-MS). The exchange of hydrogen to deuterium (HDX) of backbone amides in a protein is dependent on the presence and stability of hydrogen bonds and thus provides a sensitive probe for higher-order structure and dynamics of the target protein in solution (31, 32). HDX-MS is a non-perturbing technique that allows the collection of structural dynamics data along the entire protein backbone in a coherent manner without the need for sequence alterations or changes to the covalent structure of the protein for labeling (33). Briefly, the target protein is diluted into deuterated buffer and labeled for various time intervals. The isotopic exchange reaction is quenched by lowering pH and temperature to approximately 2.5 and 0 °C, respectively, and the protein subsequently digested using an acid-stable protease (*e.g.* pepsin). Chromatographic separation and mass analysis of these peptides, in turn, reveal the shift in mass over time (*i.e.,* deuterium uptake) of individual regions of the target protein, which is commonly referred to as local HDX analysis. A more detailed background on the HDX-MS technique and its applications in protein science can be found in several reviews (34-37).

Our HDX-MS measurements provide a detailed map of LeuT regions involved in conformational changes during substrate transport (TMs 1a/1b/2/5/6a/6b/7 and interconnecting loops IL1/EL2/EL3/EL4b) and allow for the first time an unperturbed global view on the structural dynamics associated with the outward-to-inward transition of the wild-type transporter in solution. Of special interest, our acquired HDX data suggest that several helices (TMs 1a/5/6/7 and EL4b) are partially unwound in the course of substrate transport and that these unfolding events are dynamically coupled between individual helices that form the substrate binding site and the cytoplasmic gate. Addition of Na^+^ or the combination of Na^+^ and leucine destabilized discrete structural motifs on the extracellular side, stabilized the inner gate of LeuT, and substantially reduced the rate of unfolding in individual TM helices relative to the K^+^-bound state. We envisage that partial unwinding of TM helices accompanies the outward-to-inward isomerization in LeuT and that the same concept might be relevant to related transporters bearing the LeuT fold, hence extending the general model of NSS transport mechanism. Moreover, we provide additional experimental evidence for a potential role of K^+^ in the transport cycle as K^+^ selectively shifted the conformational equilibrium of LeuT in a dose-dependent manner towards an inward-facing state under physiologically relevant concentrations.

## RESULTS

### Pinpointing LeuT Segments Involved in Substrate Transport

For the HDX experiments, we used LeuT expressed in *E. coli* C41 strain, purified and solubilized in 0.05% dodecyl-β-D-maltoside (DDM). The activity of LeuT was assessed by scintillation proximity assay (SPA) as described previously (38). Affinity for [^3^H]leucine and Na^+^-dependency were in agreement with previously published data for LeuT (38) (Supplementary Fig. 1). To examine the molecular mechanism underlying transport function of LeuT, we tracked changes in deuterium uptake as a function of time and substrate/ion composition by local HDX-MS analyses. We reasoned that transporter segments, in which the protein backbone either becomes more dynamic or exhibits structural stabilization due to the isomerization of LeuT, would display increased and decreased deuterium uptake relative to a defined reference state, respectively. Online pepsin proteolysis yielded a total of 67 LeuT peptides covering 71% of the protein sequence (Fig. 1a) that complied with the requirements to monitor the time-resolved HDX (*i.e.,* 0.25-60 min) across different functional states of the transporter (Fig. 1b).

**Figure 1.**
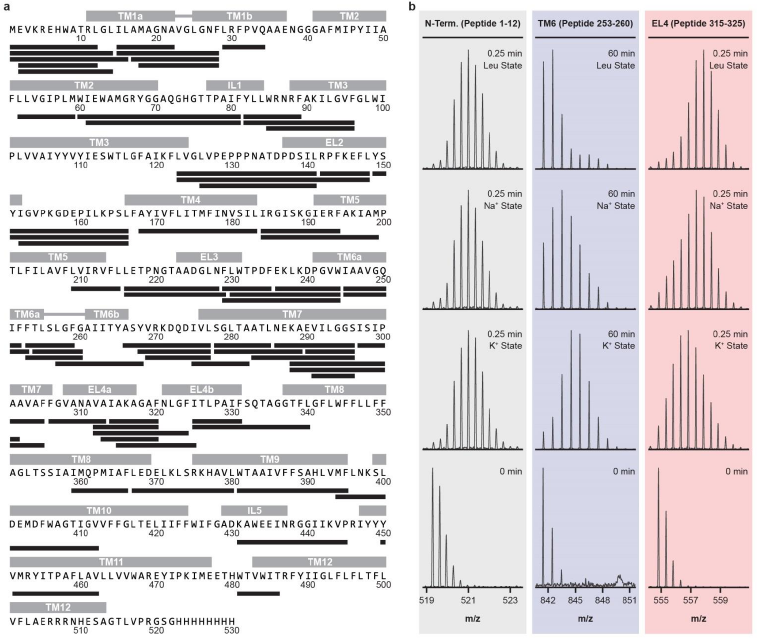
Measuring the HDX in distinct LeuT regions by mass spectrometry. **(a)** Online pepsin proteolysis yielded a total of 67 LeuT peptides suitable for local HDX-MS analysis. The identified peptides are depicted as black bars and are aligned with the corresponding LeuT sequence. The peptides cover 71% of the protein sequence. Individual structural motifs in LeuT are indicated above the protein sequence in grey color. **(b)** Representative mass spectra for individual LeuT peptides that cover the N-terminus (peptide 1-12), the substrate binding site in TM 6 (peptide 253-260), and EL4 (peptide 315-325) are shown. The isotopic envelopes shift to higher values on the m/z scale as a function of time (0.25-60 min) and ion/substrate composition (*i.e.,* K^+^, Na^+^, and Leu state) due to deuterium incorporation at the backbone amide position. Addition of Na^+^ or the combination of Na^+^ and leucine led to decreased HDX (*i.e.,* structural stabilization) in the substrate binding site in TM 6 (peptide 253-260, blue panel) and to increased HDX (*i.e.,* structural destabilization) in EL4 (peptide 315-325, red panel) relative to the K^+^ state. The exchange rate in the N-terminus (peptide 1-12, grey panel) was not affected upon addition of Na^+^ and leucine.

We chose the purification conditions (200 mM KCl, 20 mM Tris-HCl, pH 8.0, and 0.05% DDM) for the LeuT reference state and selectively shifted the conformational equilibrium of the transporter from a presumably more inward-facing open configuration (38) towards outward-oriented states of the transport cycle by varying the ion/substrate composition in a similar manner as described previously (39-41). We thus measured the local HDX in LeuT as a function of time in K^+^ (200 mM KCl, K^+^ state), Na^+^ (200 mM NaCl, Na^+^ state), and in the presence of 200 mM NaCl and varying concentrations of leucine (Leu state, *cf.* Supplementary Materials and Methods).

We first studied the deuterium uptake of LeuT in the K^+^ state. The earliest measured time-point most accurately samples the HDX of fast-exchanging, non-hydrogen bonded amide hydrogens (31) and the observed deuterium uptake after 0.25 min of labeling correlated well with the expected higher-order structure of LeuT. We observed faster exchange rates in segments encompassing the more flexible loops (Supplementary Fig. 2).

In particular, EL4a exchanged rapidly demonstrating a pronounced dynamic behavior. In contrast, most peptides that cover individual TM helices showed limited deuterium incorporation consistent with the respective backbone amide hydrogens being engaged in intramolecular hydrogen bonds. One exception is TM 4, which had an exchange rate similar to most loops. Strikingly, several LeuT peptides covering the intracellular halves of TMs 1a/5/7, the substrate binding site in TM 6, and EL4b displayed bimodal isotopic envelopes upon deuteration demonstrating that these transporter regions coexist in a folded and an unfolded state, which interconvert in solution *via* slow and correlated unfolding/refolding motions. The structural interpretation of these unusual exchange kinetics (so-called EX1 kinetics) in LeuT is described in detail in a subsequent section.

HDX results for the K^+^ state then served as a baseline for determining the impact of Na^+^ and leucine binding on conformation and dynamics in LeuT. Distinct transporter segments including TMs 1a/1b/2/5/6a/6b/7 and interconnecting loops IL1/EL2/EL3/EL4b displayed perturbed HDX in the Na^+^ and leucine-bound states relative to the K^+^ state (Fig. 2a and Fig. 2b). Mapping these differences in HDX onto available LeuT crystal structures revealed that regions exhibiting either structural stabilization or increased structural dynamics upon addition of Na^+^ or Na^+^/leucine were arranged in a symmetrical manner relative to the axis of the lipid membrane (Fig. 2c and Fig. 2d). That is, structural motifs on the extracellular side became more dynamic, whereas substrate binding sites and LeuT segments on the intracellular side exhibited structural stabilization. Furthermore, differences in exchange seemed to be confined to the functionally important 4-helix bundle domain (TMs 1/2/6/7) (42) and spatially neighboring transporter regions.

**Figure 2.**
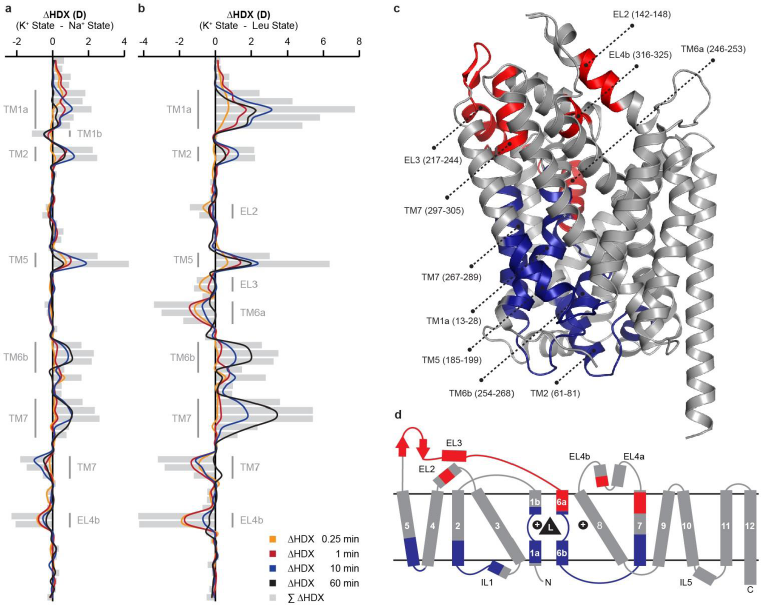
Pinpointing LeuT segments involved in substrate transport. The average relative deuterium uptake of the Na^+^ state **(a)** or the Leu state **(b)** is subtracted from the average value of the K^+^ state for each peptide and time-point. Individual LeuT peptides are plotted on the y-axis in an ordered manner starting from the N- to the C-terminus. Differences in HDX (ΔHDX, colored lines) between the Na^+^ or Leu-bound state and the K^+^ state are plotted on the x-axis. Positive and negative values indicate increased and decreased HDX in the K^+^ state, respectively. Values represent means of three independent measurements. Grey bars illustrate the sum of ΔHDX values for all sampled time-points. Structural motifs in LeuT, for which we observed Na^+^- and/or leucine-induced changes in HDX, are annotated along the y-axis. **(c)** Differences in HDX between the Leu state and the K^+^ state shown in (b) are mapped onto the LeuT crystal structure (pdb 2A65) displaying a substrate-bound, outward-facing occluded conformation. Red and blue colored regions indicate LeuT segments that became more dynamic (increased HDX) or exhibited structural stabilization (decreased HDX) in the Leu state, respectively. LeuT segments that were not impacted upon Na^+^/substrate binding or for which no HDX data could be obtained are colored in grey. The same color scheme is applied to the topology map of LeuT shown in **(d)**.

The Na^+^ and Leu states induced a similar HDX pattern along the protein backbone when analyzed relative to the results for the K^+^ state (compare Fig. 2a with Fig. 2b). This is consistent with the notion that both Na^+^ and the combination of Na^+^ and leucine shift the conformational ensemble of LeuT from an inward-facing towards an outward-facing transporter configuration. Despite perturbations in HDX being generally more pronounced for the Leu state, destabilization of EL3 and the TM 6a helix was only observed in the presence of substrate and represented an apparent dissimilarity between the Na^+^ and the Leu state. The measured HDX values for the K^+^, Na^+^, and Leu states are summarized for each peptide as deuterium uptake plots in Supplementary Figure 3. Surprisingly, except for the leucine binding site in TM 1a, varying the molar ratio between LeuT and leucine (*cf.* Supplementary Materials and Methods) did not impact the exchange rate of individual transporter segments (Supplementary Fig. 4).

### Unusual HDX Kinetics Reveal Unwinding of Helices in LeuT

Studying protein dynamics in solution by HDX-MS allows for the discovery of cooperative, local unfolding/refolding events along the protein backbone (36, 43-45), in which the residence time of the unfolded state typically exceeds hundreds of milliseconds or more (46, 47). These unfolding events are often interpreted as concerted conformational changes involving long-lived perturbations of the secondary structure within a domain. The corresponding peptide mass spectra are characterized by the time-dependent appearance of two distinct mass envelopes upon deuteration that are separated on the m/z scale. The mass envelope at lower m/z values (low-mass population) relates to the protein fraction that has not yet visited the unfolded state. Once multiple residues are simultaneously exposed to deuterated solvent through a local unfolding event, all the respective backbone amide hydrogens undergo correlated exchange before the region is able to refold (*i.e.,* the chemical exchange rate greatly exceeds the rate of refolding). This non-gradual increase in average mass, also referred to as the EX1 kinetic exchange regime, gives rise to a second mass envelope at higher m/z values (high-mass population) and directly reports on the rate of unfolding or opening (k_op_) of the affected residues. Consequently, the relative abundance of the two mass envelopes has to shift over time in favor of the high-mass population as an increasing number of protein molecules visit the unfolded transition state and become labeled. Importantly, amides exhibiting EX1 kinetics are typically engaged in a stable hydrogen-bonding network in the folded state and their exchange relies on a two-step process, in which hydrogen bonds have to be disrupted by an unfolding event prior to the isotopic exchange reaction.

Analysis of the HDX-MS data on LeuT revealed a strikingly high number of segments that exchanged according to such an EX1 kinetic regime. The domains were particularly concentrated on the intracellular side of the transporter (Fig. 3a and Fig. 3b, orange colored segments). The characteristic bimodal isotopic envelopes were consistently observed in all biological replicates (n=4) and for all studied conditions (K^+^, Na^+^, and Leu state). Importantly, in limited control experiments using wild-type LeuT embedded in phospholipid-containing bilayer nanodiscs (*cf.* Supplementary Materials and Methods), we also observed EX1-type kinetics in the same individual transporter regions (Supplementary Fig. 5). However, due to the increased sample complexity of nanodisc-embedded LeuT (*e.g.* presence of phospholipids as well as non-targeted protein species including membrane scaffold protein, MSP, and porcine pepsin), we could not achieve the same high sequence coverage as for detergent-solubilized LeuT (Supplementary Fig. 6).

**Figure 3.**
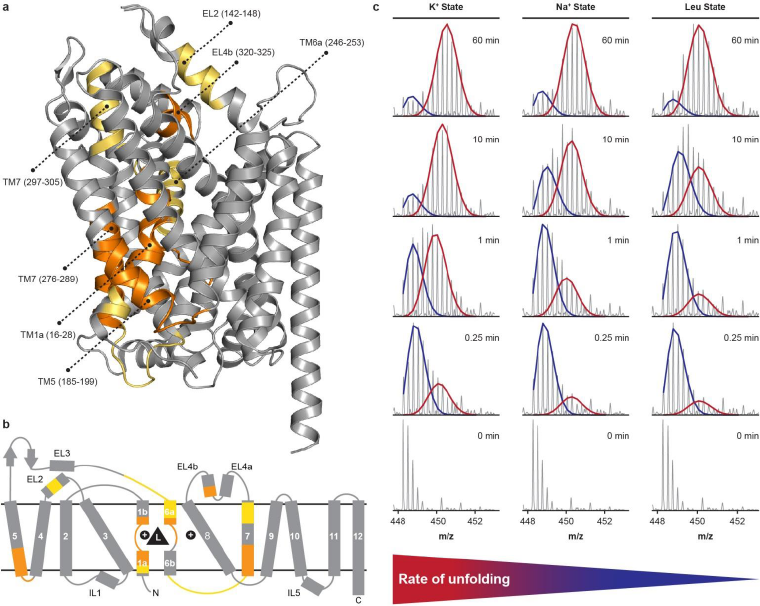
Partial unwinding of individual helices in LeuT. **(a)** LeuT segments that exchanged *via* an EX1 regime or that exhibited time-dependent broadening of the isotopic envelope upon deuteration are colored in orange and yellow in the LeuT crystal structure (pdb 2A65), respectively. The same color scheme is applied to the topology map of LeuT in **(b)**. **(c)** Representative mass spectra for peptide 184-199, which covers the intracellular half of TM 5, are shown for the K^+^, Na^+^, and Leu state and all sampled time-points. The bimodal isotopic envelopes can be fitted to a low- (blue) and high-mass (red) population. Addition of Na^+^ or the combination of Na+ and leucine substantially decreased the rate of correlated exchange, that is, the conversion of the low-mass population to the high-mass population, relative to the K^+^ state. Importantly, Na^+^- and leucine-induced stabilization of transmembrane helices was also evident for TMs 1a/6/7 with structural stabilization being consistently more pronounced in the Leu state than in the Na^+^ state (Supplementary Fig. 7).

For detergent-solubilized LeuT, we observed bimodal mass envelopes in peptides that cover the intracellular halves of TMs 1/5/7, the substrate binding site in TM 6, and EL4b. The correlated exchange in each of these structural motifs was monitored as a function of time and ion/substrate composition in at least two overlapping peptides and revealed the number of backbone amides that engage in a cooperative unfolding event. Four overlapping peptides covering residues 15-22, 15-28, 17-28, and 18-28 were used to examine the localized unfolding of TM 1. At the earliest measured time-point (0.25 min), the average difference in HDX between the low- and high-mass population in peptide 15-28 was 5.6 ± 0.2 D, which corresponds to the correlated exchange of approximately 9 backbone amide hydrogens when correcting for the measured partial loss of deuterium during HDX-MS analysis (back-exchange, *cf.* Supplementary Materials and Methods). It thereby appears that amides undergoing correlated exchange are localized in the N-terminal half of TM 1 as fewer residues underwent correlated exchange in overlapping peptides 17-28 and 18-28 (relative to peptide 15-28). This observation is further substantiated by the fact that peptides spanning residues 1-14 and 1-16 displayed distinct broadening of the isotopic envelope indicative of an EX1 exchange regime, whereas the N-terminal peptide 1-12 did not. Taken together, these data suggest that TM 1a undergoes a localized unwinding that breaks the hydrogen bonds of its backbone amides facilitating such a correlated exchange. A similar effect is observed in the intracellular half of TM 5. The local unwinding pertaining to TM 5 was studied by analyzing HDX data for peptides 184-194 and 184-199. Figure 3c shows representative mass spectra for LeuT peptide 184-199 for the K^+^, Na^+^, and Leu state and at all sampled time-points. Deconvolution of the bimodal isotopic envelopes for peptide 184-194 and 184-199 revealed the involvement of 5-6 and 7 backbone amides in a cooperative unfolding event, respectively. The increased correlated exchange in peptide 184-199 suggests that the unfolding event is centered on position 194 and neighboring residues of the TM 5 helix. In both TM 6 (peptides 253-259, 253-260, and 254-260) and EL4b (peptides 312-324 and 315-325) approximately 4-5 residues underwent correlated exchange. For peptides covering EL4b, the location of the unfolding event could be localized to residues 320-325 based on information from overlapping peptides. Finally, we observed distinct bimodal isotopic envelopes in LeuT segments corresponding to the intracellular half of TM 7, *i.e.* in three overlapping peptides covering residues 275-282, 278-285, and 278-289. The average difference in HDX between the low- and high-mass population in peptide 278-289 was 5.7 ± 0.04 D, which corresponds to the concurrent exchange of approximately 8-9 backbone amide hydrogens in the TM 7 helix. Considering that the TM 7 helix starts at residue 276, the measured correlated exchange in LeuT peptide 278-289 thus points towards a complete unwinding of the intracellular part of this transmembrane segment.

We also found distinct and time-dependent broadening of isotopic envelopes in the following LeuT segments: EL2 (peptides 123-148 and 142-148), the loop structure interconnecting EL3 and TM 6a (peptides 229-244, 230-8 244, and 236-244), TM 6a (peptides 245-252 and 245-253), the loop structure interconnecting TM 6b and TM 7 (peptides 266-277, 268-277, and 269-277), and the extracellular part of TM 7 *(e.g.* peptide 297-305). However, even though the exchange kinetics were compatible to an EX1 regime, the respective peptide mass spectra covering these individual transporter segments were not amenable to bimodal deconvolution due to various reasons such as insufficient separation of the two mass envelopes on the m/z scale or unfavorable unfolding kinetics. This is also partially illustrated in Figure 3a and 3b considering that transporter regions exhibiting broadening of the isotopic envelope (yellow colored segments) are often located proximal in sequence to segments that clearly exchanged according to an EX1 regime.

Strikingly, bimodal or broadened isotopic envelopes were solely observed in peptides covering transporter regions that were also conformationally impacted upon Na^+^ and substrate binding (compare Fig. 2c with Fig. 3a). We thus reasoned that local unfolding events, such as the unwinding of transmembrane helices, are directly linked to the substrate translocation mechanism in LeuT and potentially represent the fundamental motions that enable the transporter to isomerize between distinct functional states in an ion- and substrate-dependent manner. Addition of Na^+^ or the combination of Na^+^ and leucine substantially lowered the rate of unfolding in TMs 1a/5/6/7 relative to the K^+^ state (Fig. 3c for TM 5 peptide 184-199 and Supplementary Fig. 7), with structural stabilization of transmembrane helices being consistently more pronounced in the Leu state than in the Na^+^ state. This was in particularly apparent for peptides covering the substrate binding site in TM 6 and the intracellular half of TM 7, as binding of leucine to the transporter virtually abrogated the unwinding of these helices within the sampled time-frame (Supplementary Fig. 7b and 7c).

Based on time-resolved HDX data of regions undergoing correlated exchange (Fig. 4a), it is possible to approximate rate constants for the observed local unfolding events, that is, the rate constant of opening/unfolding (k_op_) and the half-life of the closed/folded state. We therefore plotted the relative abundance of the low-mass population in the K^+^ state for peptides amenable to bimodal deconvolution against labeling time (n=3) and fitted the data-points to an exponential decay function as described previously (47). Figure 4b shows the results of the nonlinear regression for individual LeuT peptides covering the intracellular halves of TMs 1/5/7 (peptides 15-28, 184-199, and 278-285), the substrate binding site in TM 6 (peptide 253-260), and EL4b (peptide 315-325). The curve fits correlated well with experimental data (R > 0.85) and indicated that the rate of unwinding in individual helices differs by as much as two orders of magnitude (calculated kop values ranged from 0.06 to 0.0007 s^-1^, Fig. 4c). We further noted that the determined kinetic parameters for the spatially neighboring TMs 1a and 5 were highly similar suggesting the possibility that unwinding events in these LeuT helices are dynamically coupled.

**Figure 4.**
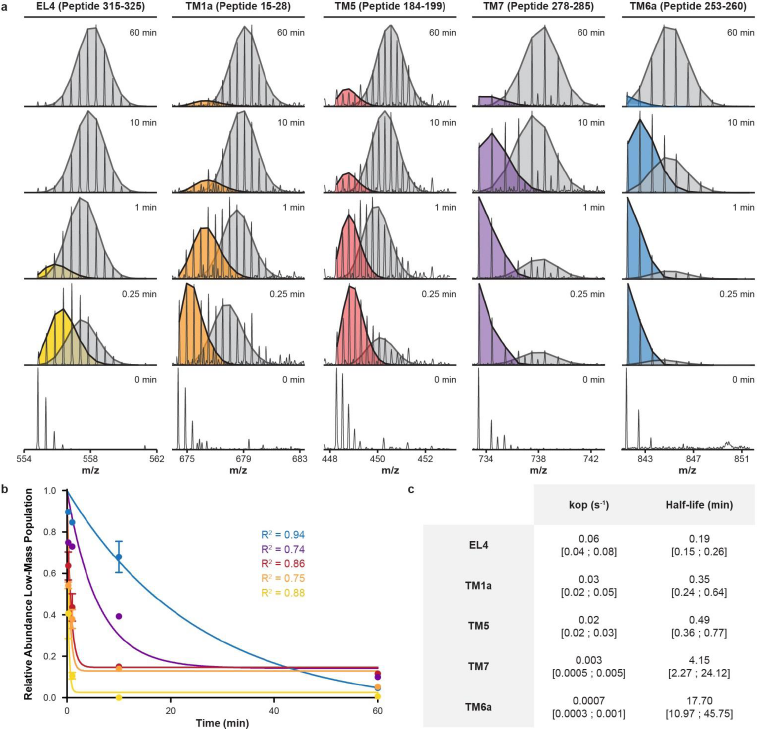
Extraction of kinetic measures for individual unfolding events in LeuT. **(a)** Time-resolved HDX data (K^+^ state) for different LeuT peptides covering EL4, TMs 1a/5/7, and the substrate binding site in TM 6 are presented. The bimodal isotopic envelopes can be fitted to a low- (colored) and high-mass (grey) population. **(b)** The average relative abundance of the low-mass population is plotted against labeling time for each peptide shown in (a). Values represent means ± standard deviation of three independent measurements. The data-points for each peptide are fitted to an exponential decay function (*cf.* Supplementary Materials and Methods). Note that each peptide is color-coded as specified in (a). **(c)** Tabular overview of the calculated kinetic parameters (*i.e.,* k_op_ and the half-life of the low-mass population) for each LeuT peptide. The best-fit values are reported together with the values defining the 95% confidence interval (in square brackets). Notably, the calculated k_op_ values for individual unfolding events in LeuT differed by as much as two orders of magnitude.

### Impact of Potassium on the Conformational Ensemble of LeuT

Recent experimental evidence (38) supports a critical role of K^+^ in the LeuT substrate transport mechanism. That is, K^+^ was shown to bind to LeuT and to promote an outward-closed/inward-facing configuration thereby inhibiting the Na^+^-dependent binding of substrate. This potentially facilitates transport by inhibiting substrate rebinding during the return step of the transporter. We therefore determined the local HDX in LeuT as a function of time and K^+^ concentration. As negative control, we used Cs^+^ (200 mM CsCl) to achieve similar ionic strength during labeling in the absence of K^+^. According to our previous data (38), Cs^+^ should not interact with LeuT under these conditions and thus allow assessment of the apo state compared to the K^+^-bound configuration. HDX-MS analysis was conducted in an identical manner as for the K^+^ and Na^+^/leucine-bound states (*cf.* Supplementary Materials and Methods) and comprised the following conditions: Cs^+^ state (200 mM CsCl), K^+^ state (200 mM KCl), and 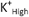 state (800 mM KCl).

Based on HDX-MS analysis, LeuT appeared to assume an outward-facing open conformation in the presence of 200 mM CsCl, which is similar to the Na^+^-bound configuration (compare Fig. 2a with Fig. 5a). We compared obtained HDX results for LeuT in Cs^+^ with the K^+^ state (Fig. 5a) in order to assess how K^+^ impacted the conformational equilibrium of LeuT. We observed perturbations in HDX for individual LeuT peptides that cover TMs 1a/5/7, the substrate binding site in TM 6, EL2, and EL4 (Fig. 5a). Intriguingly, structural motifs on the extracellular side became less dynamic (EL2 and EL4) in the presence of K^+^, whereas LeuT segments on the intracellular side (TM 1a and parts of TMs 5 and 7) exhibited structural destabilization. These changes in HDX clearly indicate a K^+^-dependent closure of the transporter to the extracellular environment. Increasing the K^+^ concentration to 800 mM appeared to cause a further shift of the LeuT conformational ensemble towards an outward-closed/inward-facing configuration (Fig. 5b). Taken together, these HDX results for LeuT in the Cs^+^, K^+^, and 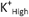 state overall support the notion of a potential role of K^+^ in the transport cycle as K^+^ seemed to selectively shift the solution-phase conformational ensemble of LeuT in a dose-dependent manner towards an outward-closed/inward-facing configuration.

**Figure 5.**
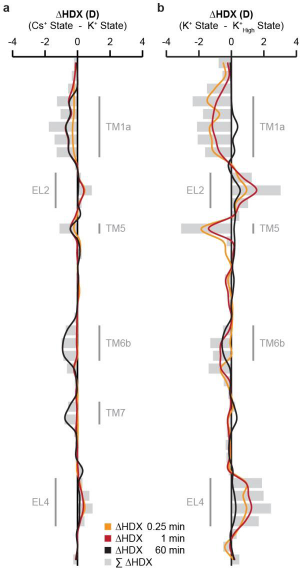
Impact of K^+^ on the conformational ensemble of LeuT. **(a)** The average relative deuterium uptake of the K^+^ state is subtracted from the average value of the Cs^+^ state for each peptide and time-point. Individual peptides are plotted on the y-axis in an ordered manner starting from the N- to the C-terminus. Differences in HDX (ΔHDX, colored lines) between the two states are plotted on the x-axis. Positive and negative values indicate increased and decreased HDX in the Cs^+^ state, respectively. Values represent means of three independent measurements. Grey bars illustrate the sum of ΔHDX values for all sampled time-points. Structural motifs in LeuT, for which we observed K^+^-induced changes in HDX, are annotated along the y-axis. **(b)** Same representation as shown in (a) for the comparison of the K^+^ and 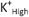 states. Positive and negative values indicate increased and decreased HDX in the K^+^ state, respectively. Values represent means of three independent measurements. Notably, (a) and (b) are consistent with a concentration-dependent, K^+^-induced closure of LeuT to the extracellular environment.

## DISCUSSION

Continuing crystallization efforts on LeuT (10-13) and other eukaryotic members of the NSS family (17, 18) have resulted in much needed insight on the structural architecture of this important class of transporters. It has become clear that NSS proteins of evolutionary distant species share a common structural fold and potentially operate *via* a preserved mechanism of action (8). LeuT belongs to the best characterized secondary active transporters with more than 50 solved crystal structures depicting distinct functional states. However, despite a plethora of LeuT crystal structures, identified key conformational states of this transporter have not yet fully depicted the structural transitions required for substrate transport (8). For instance, it has remained unclear whether isomerization of LeuT to the inward-facing open conformation is primarily driven by a rotation of the 4-helix bundle relative to the scaffold domain (TMs 3-5 and 8-10) with otherwise little additional conformational change (42) or rather relies on a more sophisticated helix bending mechanism (11). Studies taking advantage of computational and biophysical approaches have thus begun to complement the structural framework provided by X-ray crystallography to provide a better mechanistic understanding of how NSS proteins function on a molecular level by investigating the conformational dynamics of related transporters in solution. Biophysical techniques such as electron paramagnetic resonance (EPR) spectroscopy (40, 41) as well as various FRET-based strategies (23, 38, 39, 48, 49) have been applied to LeuT and demonstrated the highly dynamic nature of the transporter under different steady state conditions. However, low-resolution biophysical techniques often require changes to the covalent structure of the target protein that might perturb its function. Furthermore, these techniques are often limited to detecting conformational changes only in specific regions of the protein. Here, we have probed the solution-phase structural dynamics of LeuT in a wild type-like background by measuring the local HDX along the entire protein backbone as a function of time and ion/substrate composition.

As evident from Figure 2c, addition of Na^+^ or the combination of Na^+^ and leucine stabilized the intracellular gating region, which implies a reorientation of LeuT towards a more outward-facing configuration consistent with previous findings from both EPR (40, 41) and FRET spectroscopy studies (23, 38, 39, 48, 49). Opening of LeuT to the extracellular environment was mainly facilitated by structural and dynamical changes in the 4-helix bundle and neighboring transporter regions, clearly emphasizing the central role of the bundle domain in substrate transport. In the context of the alternating access model, it furthermore appears that individual TM helices *(e.g.* TMs 1/6/7) underwent hinge-like movements with pivot points located in the corresponding midsections. The increased differences in HDX (ΔHDX) upon leucine binding relative to the Na^+^-bound state (compare Fig. 2a with Fig. 2b) suggest that LeuT is trapped in a thermodynamically favorable outward-oriented configuration under these solution-phase conditions. Thus, the slowed exchange rates of intracellular structural motifs *(e.g.* TM 1a and TM 5) in the Na^+^/leucine-bound state reflect the increased energy barriers associated with the structural rearrangements that facilitate isotopic exchange in the EX1 regime. Because LeuT predominantly adopts an inward-facing conformation in the K^+^ reference state (40), we envisaged finding a HDX-related, common denominator between the outward-oriented Na^+^ and Leu states, which would allow us to pinpoint LeuT segments involved in the outward-to-inward transition. Based on the crystal structure of LeuT in an inward-facing open conformation, it was inferred that inward-opening in LeuT requires substantial structural rearrangements including TMs 1/2/5/6/7 and EL4b (11). Comparison of the HDX-MS results obtained for the Na^+^ state (Fig. 2a) and Leu state (Fig. 2b) supports this notion. In fact, TMs 1a/2/5/6b/7 and EL4b were the only regions in LeuT displaying a consistent trend (*i.e.,* Na^+^- and leucine-induced stabilization or destabilization of the protein backbone) for both conditions when compared to the K^+^ reference state. Strikingly, the same transporter regions, with the exception of TM 2, exchanged according to an EX1 regime signifying that the intracellular halves of TMs 1/5/7, the substrate binding site in TM 6, and EL4b exhibit unusually slow unfolding/refolding motions in solution. Interestingly, distinct structural motifs in NSS proteins have previously been suggested to undergo ion and substrate-dependent changes in secondary structure. For instance, partial unwinding of the TM 5 helix has been reported for the multi-hydrophobic amino acid transporter, MhsT, a prokaryotic member of the NSS family from *Bacillus halodurans.* X-ray crystal structures of MhsT in an occluded inward-facing state (16) suggest that a conserved GlyX_9_Pro helix-breaking motif in TM 5 (equivalent to residues Gly 190 to Pro 200 in LeuT) enables the formation of a solvent pathway for intracellular release of the Na2 ion. Moreover, it has been shown that the higher-order structure in EL4 changes upon binding of monovalent metal ions to LeuT (38).

For a more detailed structural perspective, we mapped the five segments undergoing EX1 exchange (see Fig. 3) onto three crystal structures representing markedly different states in the LeuT transport cycle: a ligand-bound, outward-occluded state (pdb 2A65) (10), an apo inward-open state (pdb 3TT3) (11), and an apo outward-open return state (pdb 5JAE) (12). Inspection of these structures (Supplementary Fig. 8) suggested the implication of most segments in the transport mechanism, either because they exhibited marked conformational change (EL4b, TM 1a, and TM 5) as quantified by RMSD measurements in Supplementary Table 1, or because they were directly in contact with the substrate-binding site (TM 1a and TM 6a). For each segment, counting the number of hydrogen bonds lost or modified in the two most different structures indicated to which degree the known crystallographic record can account for the extent of unfolding revealed by EX1 exchange (Supplementary Table 2). The largest variation was found in TM 5 segment 185-199 due to the obvious helix unwinding in structure 3TT3, and the number of hydrogen bonds lost was concomitant with the number of amide groups involved in EX1 exchange. However, it may be argued that the TM 5 conformation in 3TT3 may result from direct contacts with the antibody that stabilizes the inward-open state in the crystal (11), so that the solution-phase unwinding event relevant to function may in fact be more similar to that reported for MhsT (16). Importantly, for all other segments, the hydrogen bond variations among crystal structures cannot account for the magnitude of the observed EX1 exchange, which therefore provides novel evidence for concerted motions in LeuT not captured by available crystal structures. The most striking case is segment 279-289 of TM 7, in which 8-9 amide hydrogens exchange *via* the EX1 regime, demonstrating a concerted, slow unwinding of a substantial part of a TM helix not previously implicated in significant structural transitions during the transport cycle.

Further experimental evidence supporting the perception that the observed unwinding of TM helices in LeuT relates to the outward-to-inward transition of the transporter is provided by the fact that addition of Na^+^ or the combination of Na^+^ and leucine substantially decreased the unfolding rate of all corresponding TM domains (Fig. 3c and Supplementary Fig. 7). This observation is consistent with the concept that binding of Na^+^ and leucine stabilizes an inward-closed state in LeuT, thereby lowering the likelihood of intracellular gate opening (39). By pursuing a FRET-based imaging strategy, Zhao *et al.* highlighted that the intracellular gating dynamics in LeuT are dampened in the presence of Na^+^ (~7-fold) and that binding of leucine potentiates this effect (48). We observed a highly similar tendency with regard to the Na^+^- and substrate-induced stabilization of TM helices. Binding of leucine to LeuT virtually abrogated the unwinding of TMs 6 and 7 within the sampled timeframe (Supplementary Fig. 7b and 7c) clearly demonstrating the ability of leucine to stabilize the Na^+^-bound complex in the absence of a Na^+^ concentration gradient.

We proceeded to extract relevant kinetic measures for individual unfolding reactions observed in TMs 1/5/6/7 and EL4 (Fig. 4). We thus studied and established the spatiotemporal relationship between individual unfolding events in LeuT. Most importantly, the calculated k_op_ values did not only indicate that isomerization of LeuT to an inward-facing state relies on an ordered series of local conformational changes as suggested previously (49), but they also suggest a sequence of occurrence in the substrate translocation mechanism not accounted for in current models of NSS transport function. Based on the presented HDX data and previously published research findings, we hypothesize that substrate transport in LeuT occurs according to the following model (Fig. 6): LeuT predominantly adopts an outward-facing open conformation under apo state conditions. Obtained HDX data for LeuT in the Cs+ state and recent FRET measurements (38) support this view. The S1 binding pocket is accessible to the extracellular aqueous solution such that Na^+^ ions can bind to the Na1 and Na2 site. Of note, we observe, in agreement with other studies (39, 48), that the outward-facing open conformation is further stabilized upon formation of the Na^+^-bound complex. Subsequent binding of a substrate prompts closure of the extracellular gate giving rise to an outward-facing occluded LeuT conformation. Structural rearrangements in EL4 (calculated k_op_: 0.06 s^-1^) ensure that this loop meets the optimal length requirements to act as an ‘extracellular lid’ (10) in LeuT and simultaneously promote the forward transition in the transport cycle. Computational simulations on the bacterial hydantoin transporter Mhp1 (50), a secondary active transporter bearing the LeuT fold, support this notion. We assume, based on our HDX measurements, that the substrate-bound, outward-facing occluded state represents the thermodynamically most favorable LeuT conformation in the transport cycle. However, unwinding of the intracellular half of TM 5 allows for the formation of a solvent pathway for intracellular release of the Na2 ion, in a similar manner as described for MhsT (16). Concurrent unwinding of the TM 1a helix, which is involved in the coordination of the Na2 ion through backbone interactions at Gly 20 and Val 23 (10), further facilitates Na^+^ translocation. We reason that the structural transitions for Na2 release are dynamically coupled and find that the calculated k_op_ values for TMs 1a and 5 (0.03 and 0.02 s^-1^, respectively) are in excellent agreement with the previously reported rate constant of conformational change at the inner gate (~0.02 s^-1^) (39). The highly conserved proline kink in TM 5 and the unwound region in TM 1 may act as hinges thereby preventing the propagation of structural transitions at the inner gate to the extracellular side of the transporter. Interestingly, abstraction of the Na2 ion is associated with considerable destabilization of the substrate-bound complex and potentially induces a population shift towards an inward-facing occluded configuration compatible with the LeuT state observed by site-directed fluorescence quenching spectroscopy (23). At this point in the transport cycle, LeuT is primed for intracellular gate opening and substrate release. Our HDX measurements imply that this final isomerization step involves the observed unwinding of both the intracellular half of TM 7 (calculated k_op_: 0.003 s^-1^) and the substrate binding site in TM 6 (calculated k_op_: 0.0007 s^-1^). Intriguingly, as observed in the outward-facing occluded crystal structure (pdb 2A65) (10), TM 7 shields the Na1 ion and the substrate-binding site in TM 6 from solvent access from the intracellular side. We envisage that complete unwinding of the intracellular half of TM 7 in LeuT exposes the underlying Na1 and substrate binding site and thus provides a rational approach of how water can access the core of the transporter from the intracellular side. In other words, we propose that a second solvent pathway is formed upon unwinding of the TM 7 helix mediating access to the Na1 and S1 binding site. A similar substrate release mechanism has been suggested based on crystallographic evidence for an unrelated major facilitator superfamily transporter, the secondary active phosphate transporter PiPT (51). Compared to all other regions exhibiting EX1 kinetics, we further note that the unfolding events in TM 6 and TM 7 occur at the slowest rates, implicating these structural transitions related to substrate release as the rate limiting step in the transport cycle. In excellent agreement, we find that the determined k_op_ rate constant for unfolding of the substrate binding site in TM 6 (0.0007 s^-1^) is highly similar to the substrate turnover rate of LeuT in proteoliposomes (k_cat_ ~0.0003 s^-1^) (52). We envisage that helical interruptions in TM 6 (Ser 256 to Gly 260) and TM 7 (Gly 294) (10) may allow for structural transitions in TM 6 and TM 7 at the inner gate with minimal perturbation to the higher-order structure on the extracellular side of LeuT. Finally, our HDX measurements support the suggestion that K^+^ modulates the conformational ensemble of LeuT in a dose-dependent manner by favoring a more outward-closed/inward-facing transporter conformation (38). The functional role and the underlying molecular mechanism of K^+^-induced conformational change in LeuT, however, remains to be further elucidated.

**Figure 6.**
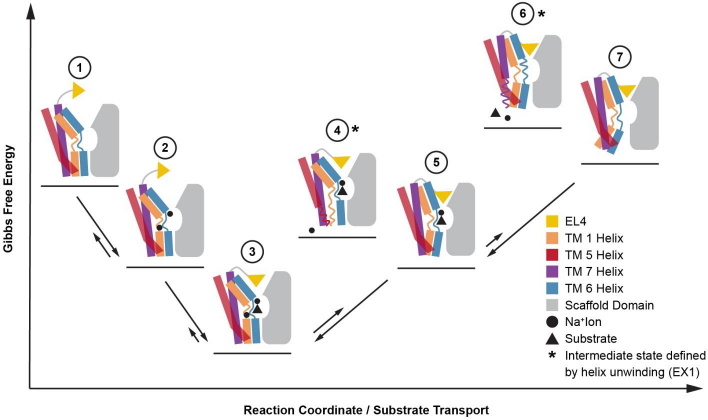
Proposed substrate transport mechanism in LeuT. **(1)** Under apo state conditions LeuT preferentially assumes an outward-facing open conformation with relatively high Gibbs free energy. **(2)** HDX is decreased upon Na^+^ binding suggesting stabilization of the protein backbone in basically the same transporter configuration as for apo state, presumably the outward-facing open. **(3)** According to crystal structures (10), binding of a transportable hydrophobic amino acid (*e.g.* leucine) prompts occlusion of the extracellular vestibule. EL4 and the water-mediated salt bridge between Arg30 and Asp404 prevent water access to the substrate binding site. Based on HDX data, it appears that the substrate-bound, outward-facing occluded state represents the thermodynamically most favorable LeuT conformation in the transport cycle. **(4)** As shown for MhsT (16) and supported by the results herein, the concurrent unwinding of the intracellular halves of TMs 1 and 5 creates a solvent pathway for intracellular release of the Na2 ion, which triggers transition of LeuT to an inward-facing occluded transporter configuration **(5)**. **(6)** The rates for the observed EX1 kinetics suggests that the release of both the substrate molecule and the Na1 ion is facilitated by a partial unwinding of the intracellular part of TM 7 thereby enabling subsequent isomerization of LeuT to an inward-facing open state **(7)**. Notably, binding of intracellular K^+^ to LeuT (not indicated in the diagram) potentially prevents substrate rebinding as well as isomerization of LeuT from an inward-facing open configuration (7) to an outward-facing open conformation (1) through a poorly understood return mechanism.

Very recently, while the current manuscript was under review, Adhikary, Deredge *et al.* have reported on the local HDX behavior of LeuT reconstituted into phospholipid-containing bilayer nanodiscs using MS (53). The authors compared the conformational differences between wild-type and a mutant LeuT (Y268A) through 35 peptic peptides covering 27% of the sequence of LeuT. The studied Y268A mutant is significant, as the mutation has been shown to induce an inward-oriented transporter configuration. Adhikary, Deredge *et al.* independently observed two regions (EL2 and intracellular region of TM 7) in the Y268A LeuT mutant to undergo concerted unfolding/refolding motions in solution. Surprisingly, the corresponding and characteristic low- and high-mass envelopes did not interconvert as a function of labeling time in the Y268A mutant. Furthermore, comparable EX1 kinetics were absent in the wild-type background. The authors inferred that this “static” bimodal behavior in the Y268A mutant signifies very slow and cooperative conformational rearrangements in the respective transporter regions. A comprehensive explanation for the lack of comparable EX1 kinetics in the wild-type transporter and how these concerted motions in EL2 and TM 7 may link to the substrate transport mechanism in LeuT was however not provided. We have independently performed a limited set of control experiments on LeuT reconstituted into phospholipid-containing nanodiscs to validate our HDX results for detergent-solubilized LeuT. We clearly observe that individual regions of nanodisc-embedded, wild-type LeuT undergo EX1 exchange with the characteristic and time-dependent interconversion of the low- and high mass envelopes (*cf.* Supplementary Fig. 5). Consequently, our mechanistic inferences based on the extensive HDX-MS measurements on detergent-solubilized LeuT (Fig. 6) appear also highly relevant when the transporter is present in a native-like phospholipid environment.

In summary, our solution-phase HDX-MS data encompass numerous experimental findings of the past decade for LeuT and other NSS-related proteins. That is, the here identified structural motifs (TMs 1a/1b/2/5/6a/6b/7 and interconnecting loops IL1/EL2/EL3/EL4b) for ion- and substrate-dependent alternating access in LeuT are identical to the ones previously proposed based on crystallographic evidence (10, 11) and biophysical measurements (23, 38-41, 48). The possibility to study the conformational dynamics of LeuT along the entire protein backbone under different steady-state conditions, however, allowed us to explore the extensive allosteric regulation in LeuT in a coherent and spatiotemporally resolved manner, which revealed novel mechanistic features inherent to the substrate transport mechanism. We propose that isomerization of LeuT to an inward-facing state relies on the partial unwinding of TMs 1a/5/7 and the substrate binding site in TM 6. Furthermore, we reason that the release of the substrate molecule and both Na^+^ ions is accomplished by these conformational rearrangements in TMs 1/5/6a/7, resulting in solvent pathways to the core of the transporter. Importantly, although our study provides direct experimental evidence for partial unwinding of TM helices in LeuT, the proposed transport model is speculative. In Figure 6, we provide a rational work hypothesis for the transport mechanism of LeuT by combining our findings from solution-phase HDX measurements with the established structural and functional framework for LeuT and related transporters. Considering that the proposed substrate transport mechanism for LeuT is facilitated by its higher-order structure, we envisage that other transporter bearing the conserved LeuT fold may operate *via* a similar mechanism of action. Finally, we are convinced that the HDX-MS technique represents a valuable approach to probe the conformation and structural dynamics of comparable transporters of similar complexity. The technique appears to hold an unrecognized potential for identification and characterization of the fundamental motions that appear to govern ion- and substrate-dependent alternating access in secondary active transporters.

## MATERIALS AND METHODS

### Chemicals and Reagents

All chemicals and reagents were purchased from Sigma Aldrich and were of the highest grade commercially available unless stated otherwise.

### Expression and Purification of LeuT

Wild-type LeuT was overexpressed in *E. coli* C41(DE3) transformed with pET16b vector encoding C-terminally His-tagged protein (expression plasmid was kindly provided by Dr. E. Gouaux, Vollum Institute, Portland, Oregon, USA). Following membrane isolation and solubilization in 1% (w/v) DDM, LeuT was purified in Buffer A (20 mM Tris-HCl, pH 8.0, 200 mM KCl, 0.05% (w/v) DDM, and 20% (v/v) glycerol) using nickel affinity chromatography, essentially as described previously (38). A brief description of the experimental procedure can be found in the Supplementary Materials and Methods.

### Expression and Purification of Membrane Scaffold Protein MSP1D1

MSP1D1 comprising a polyhistidine tag was overexpressed in *E. coli* BL21(DE3) cells and purified essentially as described previously (54). A brief description of the experimental procedure can be found in the Supplementary Materials and Methods.

### Scintillation Proximity Assay

The activity of purified LeuT was tested by assessing its binding affinity for [^3^H]leucine and Na^+^-dependency by SPA as described previously (38). A brief description of the experimental procedure can be found in the Supplementary Materials and Methods.

### Hydrogen/Deuterium Exchange

We induced different functional states of LeuT by varying the ion and substrate composition in the sample solution in a similar manner as described previously (39-41). Except for the Leu state at non-saturating conditions, different functional states were induced by adding an excess of the respective ion and substrate and the sample solution equilibrated for at least 30 min at 25 °C prior to the labeling with deuterium oxide. The concentration of Tris-HCl (20 mM, pH 8.0) and DDM (0.05% (w/v)) was kept constant for all samples throughout the labeling workflow. The corresponding sample preparation is described in more detail in the Supplementary Materials and Methods. The sample preparation procedure for nanodisc-embedded LeuT is entirely described in the Supplementary Materials and Methods.

Subsequent to the equilibration with ion and substrate, HDX was initiated by diluting LeuT 10-fold in deuterated buffer and the samples labeled at a constant temperature of 25 °C for the indicated time intervals (*i.e.,* 0.25-60 min). All labeling buffers contained 20 mM Tris-HCl, pH 8.0 and 0.05% (w/v) DDM. The labeling buffers were supplemented with the corresponding salt needed to maintain equilibration conditions. Ice-cold quench buffer (220 mM phosphate buffer, pH 2.3, and 6 M urea) was added in equal volume to the sample solution at the indicated time-points to inhibit the isotopic exchange reaction. Quenched protein samples were immediately frozen and stored at -80 °C until further use. LeuT samples, in which the ion and substrate composition matched the purification conditions and in which LeuT was labeled for 72 hours served as 90% deuterated controls. We thereby assumed that LeuT is fully labeled after the prolonged labeling period.

### Liquid Chromatography and Mass Spectrometry

To obtain local HDX information, approximately 80 pmol of labeled LeuT was loaded onto a refrigerated (0 °C) UPLC chromatographic system (nanoACQUITY-HDX technology, Waters) coupled to a hybrid Q-ToF Synapt G2 mass spectrometer (Waters, Milford, USA). Online proteolysis was accomplished at 20 °C by using an in-house packed pepsin column (IDEX, Oak Harbor, USA) containing immobilized pepsin on agarose resin beads (Thermo Scientific Pierce, Rockford, USA) and the resulting peptide mixture separated by reversed-phase liquid chromatography (see Supplementary Materials and Methods for details). The Synapt G2 mass spectrometer was interfaced with an ESI source and operated in positive ionization mode. Human Glu-Fibrinopeptide B (Sigma-Aldrich) served as an internal standard and was acquired throughout the analysis. To minimize spectral overlap, ions were further separated in the gas-phase using ion mobility. The ion mobility cell was operated at a 600 m/s wave velocity, a 40 V wave height, and a constant 90 ml/min nitrogen gas flow. Peptides were identified by data-independent MS/MS analysis (MS^E^) of LeuT (approximately 100 pmol/injection) with the ion mobility cell turned off using collision-induced dissociation and subsequent database searching in the PLGS 3.0 software.

### Peptide Identification Criteria and HDX Data Analysis

Peptide hits were filtered according to fragmentation quality (minimum fragmentation products per amino acid: 0.2), mass accuracy (maximum MH+ error: 10 ppm), and reproducibility (peptide identification in 50% of MS^E^ runs) prior to their integration into HDX analysis. HDX-MS data was processed in the DynamX 3.0 software and all peptide assignments manually verified. Noisy and overlapping HDX data was discarded from HDX analysis. Back-exchange (BE) was calculated for each peptide based on average values of at least three independent measurements according to the following equation,

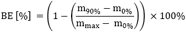

where m_90%_ is the measured average mass of the peptide for the 90% control, m_0%_ the average mass of the unlabeled peptide, and m_max_ the theoretical average mass of the fully labeled peptide. We considered that the N-terminal residue and all proline residues in a given peptide do not contribute to the measured deuterium uptake. The overall average deuterium back-exchange level in the local HDX-MS setup was calculated to be 38 ± 9%. We thereby only considered LeuT peptides that did not display a measurable difference in HDX between the 60 min time-point and the 90% control and thus appeared to be fully labeled within the sampled timeframe.

Peptides exhibiting bimodal isotopic envelopes upon deuteration were further analyzed using HX-Express 2.0. A brief description of the analysis procedure for peptides displaying EX1 kinetics can be found in Supplementary Materials and Methods.

## ACKNOWLEDGEMENTS

We would like to thank J.S. Mortensen and L. Rosenquist from the Molecular Neuropharmacology and Genetics Laboratory, University of Copenhagen, for the kind help with the SPA measurements and the purification of LeuT. We would also like to thank I.R. Möller from the Protein Analysis Group, University of Copenhagen, for fruitful scientific discussions. This work was made possible through support from the Danish Council for Independent Research (0602-02740B to KDR, 0602-02100B and 4183-00581 to CJL) and by the UCPH bioSYNergy center of excellence (CJL).

## AUTHOR CONTRIBUTIONS

KDR and CJL conceived and supervised the work. PSM and KG produced recombinant LeuT. PSM performed the HDX-MS and SPA experiments. PSM and KDR analyzed the HDX-MS data. CJL, KDR, and UG contributed with equipment, reagents, and knowhow. PSM, CJL, and KDR prepared the manuscript and all authors read and commented it.

## SUPPLEMENTARY MATERIALS AND METHODS

### Purification of LeuT

Briefly, detergent-solubilized LeuT was immobilized on Chelating Sepharose Fast Flow resin (GE Healthcare), extensively washed with ice-cold Buffer A containing step imidazole gradient (60-100 mM), and eluted in the same buffer supplemented with 300 mM imidazole. Sample purity was evaluated by SDS-PAGE analysis and protein concentration was determined by measuring the UV_280_ absorbance (ε = 1.91 cm^2^/ml). LeuT fractions with the highest protein concentrations were pooled and dialyzed against imidazole-free Buffer A for at least 16 h using 0.5 ml Slide-A-Lyzer MINI dialysis devices (10 kDa MWCO; Thermo Fisher Scientific). Dialyzed LeuT samples were stored as small aliquots at -80 °C until further use.

### Purification of MSP1D1

MSP1D1 was isolated by nickel affinity chromatography and sample purity evaluated by SDS-PAGE analysis. The polyhistidine tag was removed by overnight incubation of purified MSP1D1 in the presence of 3% (w/w) tobacco etch virus protease under constant rotation at room temperature. Cleaved MSP1D1 protein was collected using nickel affinity chromatography and dialyzed against buffer containing 20 mM Tris-HCl, pH 8.0, and 100 mM NaCl. Following up-concentration of MSP1D1 to 4 mg/ml using spin filter units (Vivaspin centrifugal concentrator, 10 kDa MWCO, Sartorius Biotech), sample aliquots were stored at -80 °C until further use.

### Scintillation Proximity Assay

Briefly, for saturation binding, 0.5 μg/ml of purified LeuT was mixed with 0.125 mg/ml YSi-Cu His-Tag SPA beads (PerkinElmer) and varying concentrations of [^3^H]leucine (PerkinElmer, 10.8 Ci/mmol), all diluted in binding buffer (20 mM Tris-HCl, pH 8.0, 200 mM NaCl, 0.05% (w/v) DDM). Na^+^-dependency was determined using fixed 100 nM [^3^H]leucine, in assay buffer supplemented with varying concentrations of NaCl while ionic strength was maintained with KCl. Nonspecific background was determined in the presence of unlabeled leucine. Binding of [^3^H]leucine was monitored employing MicroBeta scintillation counter (PerkinElmer) and data were fitted using the GraphPad Prism 6 software to a one-site saturation or dose-response function to obtain K_d_ and EC_50_ values, respectively.

### Hydrogen/Deuterium Exchange of Detergent-Solubilized LeuT

To obtain the Na^+^ and Cs^+^ states, purified LeuT was dialyzed overnight using 0.5 ml Slide-A-Lyzer MINI dialysis devices (10 kDa MWCO) at 4 °C against modified Buffer A containing 20 mM Tris-HCl, pH 8.0, 0.05% (w/v) DDM, 20% (v/v) glycerol, and either 200 mM NaCl or 200 mM CsCl. The 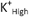 state was obtained by manually adding a stock solution containing 20 mM Tris-HCl, pH 8.0, 0.05% (w/v) DDM, and 3.2 M KCl to the purified LeuT stock to reach a final KCl concentration of 800 mM. Two different Leu states were prepared and analyzed in the present study. First, we studied the HDX of LeuT in presence of 200 mM NaCl and a non-saturating leucine concentration. For this reason, NaCl was manually added to the purified LeuT stock to a final concentration of 200 mM, whereas leucine was present in the deuterated buffer. The leucine concentration during the labeling reaction was 1 μM, which resulted in approximately 60% transporter occupancy. Note that LeuT was not equilibrated with the substrate leucine prior to the labeling reaction. Second, we prepared samples, in which LeuT was equilibrated with a 3-fold molar excess of leucine prior to deuteration. Thereby, NaCl and leucine were manually added to the purified LeuT stock to a final concentration of 200 mM and 50 μM, respectively, which resulted in >99% transporter occupancy.

### Reconstitution of LeuT into Phospholipid-Containing Bilayer Nanodiscs

Preparation of phospholipids and nanodisc assembly were performed in a similar manner as described previously (54). The phospholipids 1-palmitoyl-2-oleoyphosphatidylcholine (POPC) (Avanti Polar Lipids) and 1-palmitoyl-2-oleyl-sn-glycero-3-phosphoracglycerol (POPG) (Avanti Polar Lipids) were prepared in chloroform and mixed in a 3:2 molar ratio. Chloroform was evaporated under nitrogen gas for 1 h and residual solvent removed by overnight incubation under vacuum. The thin lipid film was resuspended in buffer containing 10 mM Tris-HCl, pH 7.5, 100 mM choline chloride (ChCl), and 80 mM sodium cholate to a final lipid concentration of 50 mM.

LeuT, MSP, lipids, and DDM detergent were mixed in a molar ratio of 0.1:1:80:240, respectively, and incubated on ice for 1 h. Detergent was removed by addition of 80% (w/v) Biobeads (Bio-Rad SM-2 Resin) and left for 1 h on ice prior to overnight incubation at 4 °C. Loaded nanodiscs were separated from empty nanodiscs by nickel affinity chromatography and eluted in buffer containing 20 mM Tris-HCl, pH 8.0, 200 mM KCl, and 300 mM imidazole. Loaded nanodiscs were subsequently subjected to size exclusion chromatography using a Superdex 200 10/300 GL column (GE Healthcare) pre-equilibrated with 20 mM Tris-HCl, pH 7.50, and 100 mM ChCl.

### Hydrogen/Deuterium Exchange of LeuT Reconstituted into Nanodiscs

Samples of LeuT reconstituted into nanodiscs were buffer exchanged into buffer containing 20 mM Tris-HCl, pH 8.0, and 200 mM KCl. HDX was initiated by diluting LeuT 10-fold in a matching deuterated buffer and the samples labeled at 25 °C for various time intervals (*i.e.,* 0.25-60 min). A total of 43 μl of ice-cold quench solution (0.5% formic acid and 6 M urea) was added to 50 μl of sample solution at the indicated time-points to inhibit the isotopic exchange reaction. The quenched protein samples were kept on ice for the following sample preparation to minimize back-exchange. The nanodiscs were disassembled by adding an ice-cold stock solution of sodium cholate to reach a sodium cholate:lipid ratio of 25:1, respectively. Subsequently, LeuT was subjected to offline proteolysis for 2 min on ice by adding 6 μg porcine pepsin. In the last minute of proteolysis, the quenched sample was transferred into a cooled spin filter (Costar Spin-X^®^ 2 mL centrifuge tube filter 0.45 μm cellulose acetate) containing 3 mg zirconia-coated silica particles (HybridSPE^®^-Phospholipid cartridge, Supelco). The quenched sample was centrifuged at 0 °C and 3000 g for 1 min and the flow through collected. Finally, quenched protein samples were immediately frozen and stored at -80 °C until further use.

### Liquid Chromatography

Peptic peptides were trapped on a C18 column (ACQUITY UPLC BEH C18 1.7 μm VanGuard column Waters, Milford, USA) and the sample efficiently desalted for 3 min with mobile phase A (0.23% (v/v) formic acid) at a constant flow rate of 200 μl/min. Peptides were gradually eluted by applying a 9 min linear gradient at a 40 μl/min flow rate and increasing concentrations (8% - 50% (v/v)) of mobile phase B (acetonitrile, 0.23% (v/v) formic acid). Chromatographic separation was accomplished by using a C18 guard column (ACQUITY UPLC BEH C18 1.7 μm VanGuard column Waters, Milford, USA) and a C18 analytical column (ACQUITY UPLC BEH C18 1.7 μm, 1x100 mm column Waters, Milford, USA).

### Mass Spectrometry

All HDX-MS data was obtained using the sample preparation workflows described above and in the main body of the manuscript, however data presented in Figure 5a was obtained employing a more sensitive Synapt G2-Si mass spectrometer as opposed to the Synapt G2 mass spectrometer used in all other experiments. Thus, while the two datasets shown in Figure 5a and 5b can be compared qualitatively *(i.e.,* in terms of comparing regions of the LeuT backbone, in which Cs^+^ and K^+^ induces changes in HDX), the relative magnitudes in ΔHDX might differ due to differences in the MS setup and thus a direct comparison of ΔHDX values between Figure 5a and 5b should be avoided.

### HDX Data Analysis

To determine the number of backbone amide hydrogens undergoing correlated exchange (#NHs), we first calculated the difference in HDX between the low- and high-mass population (ΔHDX, n=3) and corrected this value according to the measured back-exchange (BE, defined in Materials and Methods) of the respective peptide by using the following equation

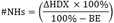

To approximate critical kinetic measures *(i.e.,* k_op_ and the half-life of the folded state) for individual unfolding events in LeuT, we first determined the relative abundance of the two isotopic envelopes for a given peptide using HX-Express 2.0. We subsequently plotted the relative abundance of the low-mass population (n=3) against labeling time (*i.e.,* 0.25-60 min) based on HDX data for LeuT in K^+^ reference state. Finally, the GraphPad Prism 6 software was used to fit the individual data-points to the following exponential decay function

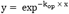

in order to obtain the best-fit values for the kinetic parameters. HDX results were mapped onto the LeuT crystal structure (pdb 2A65) using Pymol (http://pymol.sourceforge.net/).

---

**SI Table 1.** RMSD of backbone atoms of EX1 segments in X-ray structures 3TT3 and 5JAE with respect to structure 2A65 after alignment on helical regions.

| Location | Segment | 2A65 - 3TT3<br>RMSD [Å] | 2A65 - 5JAE<br>RMSD [Å] |
| --- | --- | --- | --- |
| EL4b | 320-325 | 2.7 | 1.1 |
| TM1a | 16-28 | 6.6 | 1.7 |
| TM5 | 185-199 | 5.8 | 0.4 |
| TM7 | 279-289 | 1.2 | 0.4 |
| TM6a | 254-260 | 1.8 | 1.0 |

**SI Table 2.** Hydrogen bond analysis of backbone amide groups in three X-ray structures of LeuT. These structures represent a ligand-bound, outward-occluded state (pdb 2A65), an apo inward-open state (pdb 3TT3), and an apo outward-open return state (pdb 5JAE). For each segment we show the number of deuteriums exchanged in the EX1 regime, the maximum variation in number of hydrogen bonds between any two structures, the maximum number of hydrogen bonds that have changed between any two structures (*i.e.,* the acceptor is different, or the hydrogen bond is formed/broken).

| Location | Segment | Number of<br>deuteriums<br>exchanged | Max variation<br>in HB number | Max number<br>of different<br>HBs | Most different<br>structures |
| --- | --- | --- | --- | --- | --- |
| EL4b | 320-325 | 4-5 | 1 | 4 | 3TT3 - 5JAE |
| TM1a | 16-28 | 9 | 4 | 7 | 2A65 - 3TT3 |
| TM5 | 185-199 | 7 | 8 | 9 | 2A65 - 3TT3 |
| TM7 | 279-289 | 8-9 | 0 | 0 | 2A65 - 3TT3 |
| TM6a | 254-260 | 4-5 | 0 | 2 | 3TT3 - 5JAE |

**SI Figure 1.**
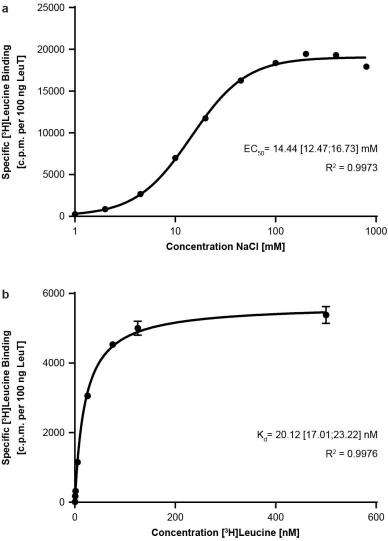
Na^+^-dependent binding of [^3^H]leucine to purified LeuT. The activity of purified LeuT was evaluated by assessing its binding affinity for [^3^H]leucine and Na^+^-dependency by SPA. **(a)** Dose-response curve for purified LeuT in the presence of fixed 100 nM [^3^H]leucine and varying concentrations of NaCl. The calculated EC_50_ value for Na^+^ was 14 [12;17] mM (mean [95% confidence interval], n=3). **(b)** One-site saturation curve for purified LeuT in the presence of 200 mM NaCl and varying concentrations of [^3^H]leucine. The calculated K_d_ for [^3^H]leucine was 20 [17;23] nM (n=2).

**SI Figure 2.**
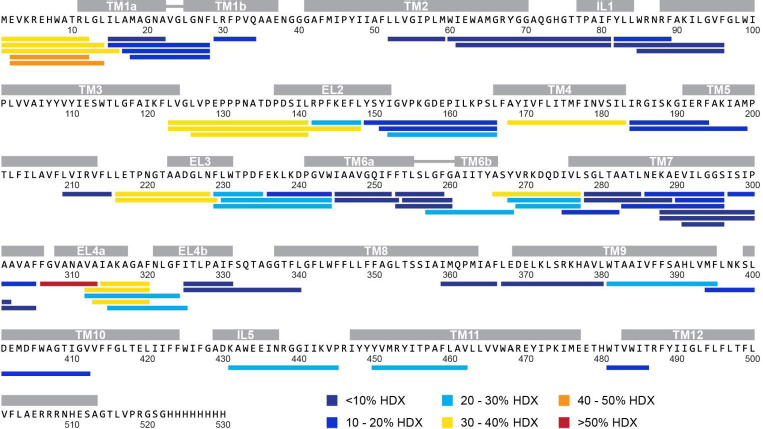
Correlation between the measured HDX and secondary structure elements in LeuT. The sequence coverage map for LeuT is shown. Individual structural motifs in LeuT are indicated above the protein sequence in grey color. Identified peptides are depicted as colored bars and are aligned with the corresponding LeuT sequence. Individual peptides are colored according to the measured deuteration level at the earliest measured time-point (0.25 min) for the K^+^ reference state (*cf.* color scheme). Deuteration levels were obtained by comparing for each peptide the measured relative deuterium uptake with a theoretical, maximal deuterium uptake value (maximal deuterium uptake = number of residues - N-terminus - number of proline residues). Only the low-mass population was considered for peptides displaying bimodal isotopic envelopes. Notably, the more flexible loop regions appeared to exchange faster than individual TM helices.

**SI Figure 3.**
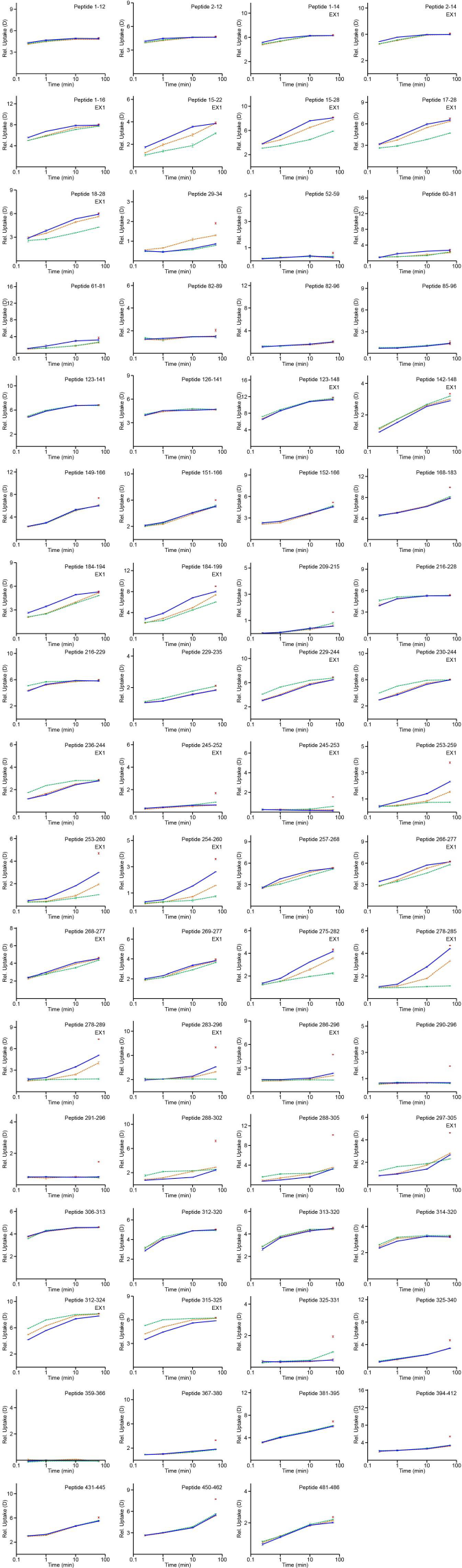
Deuterium uptake plots for detergent-solubilized LeuT. Deuterium uptake plots for all identified LeuT peptides are presented. The measured relative deuterium uptake for individual LeuT peptides is plotted against labeling time *(i.e.,* 0.25-60 min). Blue, orange, and green curves illustrate the relative deuterium uptake for detergent-solubilized LeuT in the K^+^, Na^+^, and Leu state, respectively. The red colored square at the 60 min time-point indicates the measured relative deuterium uptake for the fully labeled control sample. Values represent means of three independent measurements. The corresponding standard deviations are indicated but are in most instances too small to be visible. For peptides displaying bimodal or broadened isotopic envelopes upon deuteration (marked as ‘EX1’), we plotted an intensity-weighted, average deuterium uptake value accounting for both the low- and high-mass population. The HDX-MS data presented in this Figure is also partially shown in form of difference charts in Figure 2a and 2b.

**SI Figure 4.**
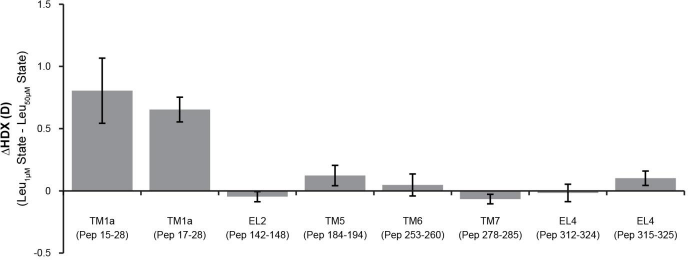
Impact of different leucine concentrations on local HDX rates in LeuT. The average relative deuterium uptake of the Leu_50μM_ state is subtracted from the average value of the Leu_1μM_ state for each peptide that is shown (*cf.* Supplementary Materials and Methods). These LeuT peptides were identified to be the most sensitive ones to Na^+^ and leucine binding (Fig 2a and 2b) and they are denoted along the x-axis. Differences in HDX (ΔHDX) between the two substrate-bound states are plotted on the y-axis. Positive and negative ΔHDX values indicate increased and decreased HDX in the Leu_1μM_ state, respectively. Values represent means and standard deviations of three independent measurements for the 0.25 min time-point. Notably, increasing the molar ratio between leucine and LeuT resulted in decreased HDX in TM 1a but did not impact the exchange rate of other structural motifs of the transporter.

**SI Figure 5.**
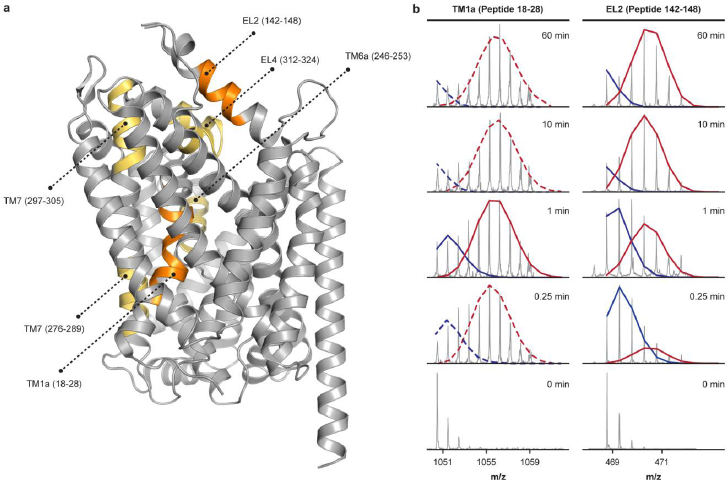
Partial unwinding of individual helices in LeuT reconstituted into phospholipid-containing bilayer nanodiscs. **(a)** LeuT segments that exchanged *via* an EX1 regime when the transporter was labeled in a phospholipidic environment are colored in orange in the LeuT crystal structure (pdb 2A65). LeuT segments for which we observed signs of EX1 kinetics but for which bimodal deconvolution could not be performed are indicated in yellow. **(b)** Representative mass spectra for individual peptides covering TM 1a and EL2 are shown for LeuT reconstituted in nanodiscs and in the presence of 200 mM K^+^. The bimodal isotopic envelopes can be fitted to a low- (blue) and high-mass (red) population. Bimodal deconvolution was accomplished using HX Express 2.0 (solid lines). In some instances, bimodal deconvolution did not result in a qualitatively meaningful fit and the depicted low- and high-mass population (dashed lines) was adjusted manually for visual guidance. Notably, we observe a time-dependent interconversion of the low- and high-mass population compatible to an EX1 exchange regime.

**SI Figure 6.**
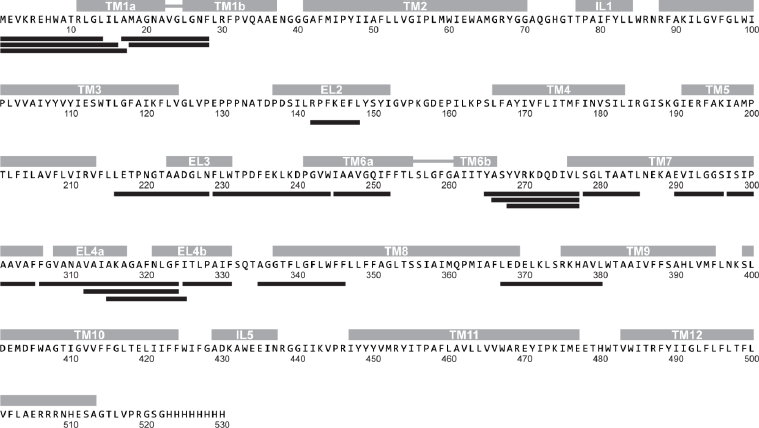
Sequence coverage map for LeuT reconstituted into phospholipid-containing bilayer nanodiscs. Offline pepsin proteolysis yielded a total of 21 peptides suitable for local HDX-MS analysis. The identified peptides are depicted as black bars and are aligned with the corresponding LeuT sequence. The peptides cover 30% of the protein sequence. Individual structural motifs in LeuT are indicated above the protein sequence in grey color. Notably, due to the increased sample complexity for nanodisc-embedded LeuT, we could not achieve the same high sequence coverage as for detergent-solubilized LeuT (*cf.* Fig. 1a).

**SI Figure 7.**
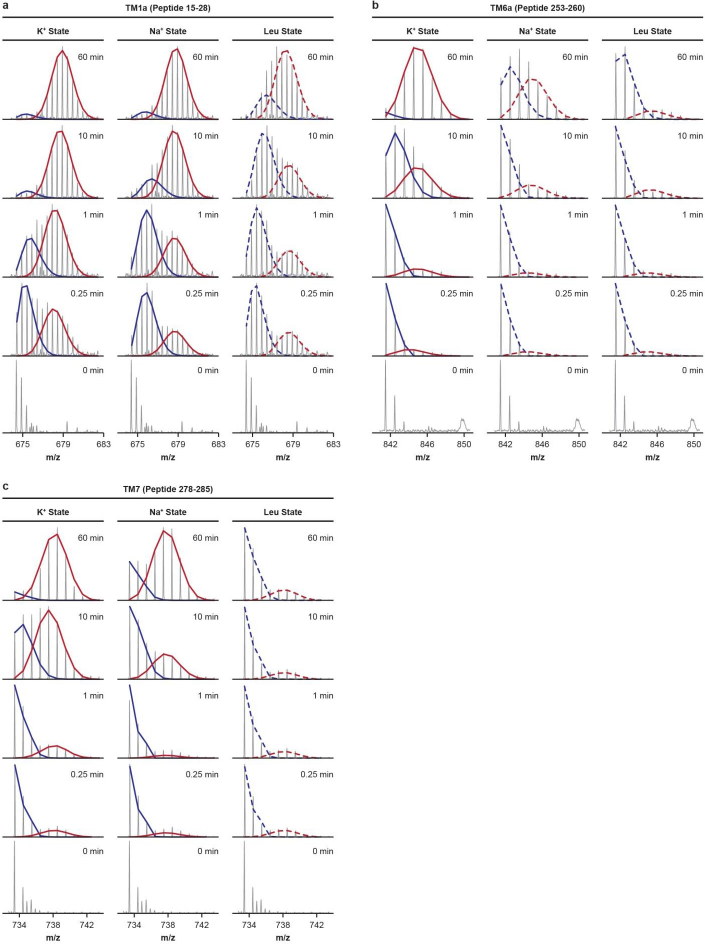
Na^+^- and substrate-induced stabilization of TM helices. Representative mass spectra for individual peptides covering **(a)** TM 1a (peptide 15-28), **(b)** the substrate binding site in TM 6 (peptide 253-260), and **(c)** the intracellular half of TM 7 (peptide 278-285) are shown for the K^+^, Na^+^, and Leu state and all sampled time-points. The bimodal isotopic envelopes can be fitted to a low- (blue) and high-mass (red) population. Bimodal deconvolution was accomplished using HX Express 2.0 (solid lines). In some instances, bimodal deconvolution did not result in a qualitatively meaningful fit and the depicted low- and high-mass population (dashed lines) was adjusted manually for visual guidance. Notably, for all peptides shown above, addition of Na^+^ or the combination of Na^+^ and leucine substantially decreased the rate of correlated exchange relative to the K^+^ state.

**SI Figure 8.**
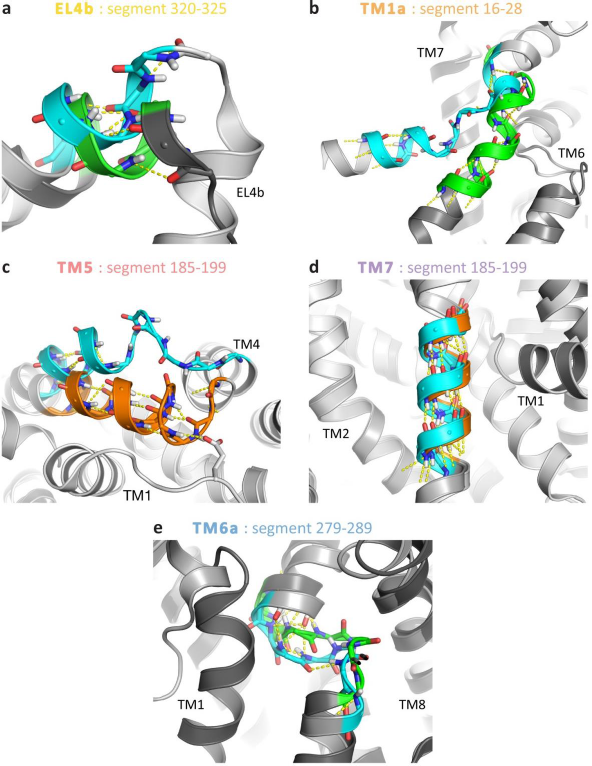
Compared conformations of EX1 segments in crystal structures. Each panel shows the two most different out of three selected X-ray structures: a ligand-bound, outward-occluded state (pdb 2A65, orange), an apo inward-open state (pdb 3TT3, cyan), and an apo outward-open return state (pdb 5JAE, green). Hydrogen bonds, defined with a donor-acceptor distance of 3.5 Å and an angle cutoff of 50°, are represented with dashed yellow lines. **(a)** EL4b segment 320-325 seen from the “inside”, *i.e.* outwards from the approximate position of the ligand; **(b)** TM 1a segment 16-28 seen from the membrane plane. TM 5 and TM 8 have been removed for clarity; **(c)** TM 5 segment 185-199 seen from the intracellular side; **(d)** TM 7 segment 279-289 seen from the membrane plane; **(e)** TM 6a segment 254-260 seen from the membrane plane. TM 5 and TM 8 have been removed for clarity.

